# A Meta-Analysis of Alzheimer’s Disease Brain Transcriptomic Data

**DOI:** 10.1101/480459

**Authors:** Hamel Patel, Richard J.B Dobson, Stephen J Newhouse

## Abstract

**Background:** Microarray technologies have identified imbalances in the expression of specific genes and biological pathways in Alzheimer’s disease (AD) brains. However, there is a lack of reproducibility across individual AD studies, and many related neurodegenerative and mental health disorders exhibit similar perturbations. We are yet to identify robust transcriptomic changes specific to AD brains.

**Methods and Results:** Twenty-two AD, eight Schizophrenia, five Bipolar Disorder, four Huntington's disease, two Major Depressive Disorder and one Parkinson’s disease dataset totalling 2667 samples and mapping to four different brain regions (Temporal lobe, Frontal lobe, Parietal lobe and Cerebellum) were analysed. Differential expression analysis was performed independently in each dataset, followed by meta-analysis using a combining p-value method known as Adaptively Weighted with One-sided Correction. This identified 323, 435, 1023 and 828 differentially expressed genes specific to the AD temporal lobe, frontal lobe, parietal lobe and cerebellum brain regions respectively. Seven of these genes were consistently perturbed across all AD brain regions with SPCS1 gene expression pattern replicating in RNA-seq data. A further nineteen genes were perturbed specifically in AD brain regions affected by both plaques and tangles, suggesting possible involvement in AD neuropathology. Biological pathways involved in the “metabolism of proteins” and viral components were significantly enriched across AD brains.

**Conclusion:** This study solely relied on publicly available microarray data, which too often lacks appropriate phenotypic information for robust data analysis and needs to be addressed by future studies. Nevertheless, with the information available, we were able to identify specific transcriptomic changes in AD brains which could make a significant contribution towards the understanding of AD disease mechanisms and may also provide new therapeutic targets.

## INTRODUCTION

Alzheimer’s disease (AD) is the most common form of dementia affecting over 44 million individuals worldwide, and numbers are expected to triple by 2050 [1]. The hallmark of the disease is characterised by the abnormal brain accumulation of amyloid-β (Aβ) protein and hyperphosphorylated tau filaments, which forms structures known as plaques and tangles respectively. The accumulation of these proteins contributes to the loss of connections between neurone synapses, leading to the loss of brain tissue and disruption of normal cognitive functions.

As AD progresses, the spread of plaques and tangles in the brain usually occurs in a predictable pattern and can begin up to 18 years prior to the onset of clinical symptoms [2]. In the earliest stages of the disease, plaques and tangles form in areas of the brain primarily involved in learning and memory, specifically the hippocampus and entorhinal cortex, both situated in the temporal lobe (TL) region [3]. Next, the frontal lobe (FL), a region involved in voluntary movement, is affected, followed by the parietal lobe (PL), a region involved in processing reading and writing. In the later stage of the disease, the occipital lobe, a region involved in processing information from the eyes, can become affected, followed by the cerebellum (CB), a region which receives information from the sensory systems and the spinal cord to regulates motor movement. Nerve cell death, tissue loss and atrophy occur throughout the brain as AD progresses, leading to the manifestation of clinical symptoms associated with loss of normal brain function. However, not all brain regions are neuropathologically affected in the same manner. The CB, which only accounts for 10% of the brain but contains over 50% of the brains total neurones, is often neglected in AD research because it is generally considered to be partially spared from the disease as plaques are only occasionally seen but tangles are generally not reported [4] [5].

The histopathological spread of the disease is well documented, and with the advent of high throughput genomics approaches, we are now able to study the transcriptomic and biological pathways disrupted in AD brains. Microarrays can simultaneously examine thousands of genes, providing an opportunity to identify imbalances in the expression of specific genes and biological pathways. However, microarray reproducibility has always been questionable, with replication of differentially expressed genes (DEG’s) very poor [6]. For example, two independent microarray transcriptomic studies performed differential expression analysis in the hippocampus of AD brains. The first study by Miller et al. identified 600 DEG’s [7], and a similar study by Hokama et al. identified 1071 DEG’s [8]. An overlap of 105 DEG’s exist between the two studies; however, after accounting for multiple testing, no gene was replicated between the two studies. The Miller study consisted of 7 AD and 10 control subjects expression profiled on the Affymetrix platform while the Hakoma study consisted of 31 AD and 32 control subjects expression profiled on the Illumina platform. Replication between the Illumina and Affymetrix platform has been shown to be generally very high [9]; therefore, the lack of replication between the two studies is probably down to a range of other factors including low statistical power, sampling bias and disease heterogeneity.

Unlike DEG’s, replication of the molecular changes at a pathway level are more consistent and have provided insights into the biological processes disturbed in AD. Numerous studies have consistently highlighted disruptions in immune response [10] [11] [12] [13], protein transcription/translation [10] [11] [14] [15] [16] [17], calcium signalling [10] [18] [19], MAPK signalling [16] [7], various metabolism pathways such as carbohydrates [16], lipids [16] [20], glucose [21] [22] [17], and iron [11] [23], chemical synapse [18] [7] [19] and neurotransmitter pathways [11] [18] [19]. However, many of these pathways have also been suggested to be disrupted in other brain-related disorders. For example, disruptions in calcium signalling, MAPK, chemical synapse and various neurotransmitter pathways have also been implicated in Parkinsons’s Disease (PD) [24] [25]. In addition, glucose metabolism, protein translation, and various neurotransmission pathways have also been suggested to be disrupted in Bipolar Disorder (BD) [26] [27] [28] [29]. Although the biological disruptions involved in AD are steadily being identified, many other neurodegenerative and mental disorders are showing similar perturbations. We are yet to identify robust transcriptomic changes specific to AD brains.

In this study, we combined publicly available microarray gene expression data generated from AD human brain tissue and matched cognitively healthy controls to conduct the most extensive AD transcriptomic microarray meta-analyses known to date. We generate AD expression profiles across the temporal lobe, frontal lobe, parietal lobe and cerebellum brain regions. We further refine each expression profile by removing perturbations seen in other neurodegenerative and mental disorders (PD, BD, Schizophrenia [SCZ], Major Depressive Disorder [MDD] and Huntington’s Disease [HD]) to decipher specific transcriptomic changes occurring in human AD brains. These AD-specific brain changes may provide new insight and a better understanding of the disease mechanism, which in turn could provide new therapeutic targets for preventing and curing AD.

## MATERIALS AND METHODS

### Selection of publicly available microarray studies

Publicly available microarray gene expression data was sourced from the Accelerating Medicines Partnership-Alzheimer’s Disease AMP-AD (doi:10.7303/syn2580853, doi:10.1038/ng.305, doi:10.1371/journal.pgen.1002707, doi:10.1038/ng.305, doi:10.1038/sdata.2016.89, doi:10.1038/sdata.2018.185) and ArrayExpress (https://www.ebi.ac.uk/arrayexpress/) in June 2016. For a study to be selected for inclusion, the data had to (1) be generated from a neurodegenerative or mental health disorder, (2) be sampled from human brain tissue, (3) have gene expression measured on either the Affymetrix or Illumina microarray platform, (4) contain both diseased and suitably matched healthy controls in the same experimental batch and (5) contain at least 10 samples from both the diseased and control group.

### Microarray gene expression data pre-processing

Data analysis was performed in RStudio (version 0.99.467) using R (version 3.2.2). All data analysis scripts used in this study are available at https://doi.org/10.5281/zenodo.823256. In brief, raw Affymetrix microarray gene expression data was “mas5” background corrected using R package “affy” (version 1.42.3) and raw Illumina microarray gene expression data Maximum Likelihood Estimation (MLE) background corrected using R package “MBCB” (version 1.18.0). Studies with samples extracted from multiple tissues were separated into tissue-specific matrices, log2 transformed and then Robust Spline Normalised (RSN) using R package “lumi” (version 2.16.0).

BRAAK staging is a measure of AD pathology and ranges from I-VI. In general, stages I-II, III-IV and V-VI represent the “low likelihood of AD”, “probable AD” and “definite AD” respectively [30]. To maintain homogeneity within the sample groups and to be able to infer pathological related genetic changes, if BRAAK staging was available, clinical AD samples with BRAAK scores ≤ 3 or clinical control samples with BRAAK scores ≥ 3 were removed from further analysis.

Gender was predicted using the R package “massiR” (version 1.0.1) and used to subset the data into four groups based on diagnosis (case/control) and gender (male/female). Next, probes below the th 90 percentile of the log2 expression scale in over 80% of samples were deemed “not reliably detected” and were excluded from further analysis to eliminate noise [31] and increase power [32].

Publicly available data is often accompanied by a lack of sample processing information, making it impossible to adjust for known systematic errors introduced when samples are processed in multiple batches, a term often known as “batch effects”. To account for both known and latent variation, batch effects were estimated and removed using the Principal Component Analysis (PCA) and Surrogate Variable Analysis (SVA) using the R package “sva” (version 3.10.0). Gender and diagnosis information were used as covariates in sva when correcting for batch effects. Outlying samples were iteratively identified and removed from each gender and diagnosis group using fundamental network concepts described in [33]. Platform-specific probe ID’s were converted to Entrez Gene ID’s using the BeadArray corresponding R annotation files (“hgu133plus2.db”, “hgu133a.db”, “hgu133b.db”, “hugene10sttranscriptcluster.db”, “illuminaHumanv4.db”, “illuminaHumanv3.db”) and differential expression analysis was performed within each dataset using the R package “limma” (version 3.20.9).

Finally, study compatibility analysis was investigated through the R package “MetaOmics” (version 0.1.13). This package uses differentially expressed genes (DEGs), co-expression and enriched biological pathways analysis to generate six quantified measures that are used to generate a PCA plot. The direction of each Quality Control (QC) measure is juxtaposed on top of the two-dimensional PC subspace using arrows. Datasets in the negative region of the arrows were classed as outliers [34] and were removed from further analysis.

### Meta-analysis

Datasets were grouped by the primary cerebral cortex lobes (TL, FL, PL) and the CB. Meta-analysis was performed using a “combining p-values” method known as “Adaptively Weighted with One-sided Correction” (AW.OC), implemented through the R package “MetaDE” (version 1.0.5)[34]. A combining p-value method was chosen to address the biases introduced from different platforms. AW.OC was chosen as it permits missing information across datasets which are introduced by combining data generated from different microarray platforms and expression chips. This avoids the need to subset individual datasets to common probes, which essentially allows for the maximum number of genes to be analysed. Furthermore, the method provides additional information on which dataset is contributing towards the meta-analysis p-value, and has been shown to be amongst the best performing meta-analysis methods for combining p-values for biological associations [35]. The meta-analysis method does not provide an overall directional change for each gene; therefore, the standard error (SE) was calculated from the DE logFC values of each gene across the AW assigned significant datasets and used for standard meta-summary estimate analysis using the R package “rmeta” (version 2.16). This served as the “meta expression” change in downstream analysis where positive values represent a gene being up-regulated in AD and negative values as being down-regulated in AD Selecting DE genes based on an arbitrary expression change significantly influences the interpretation of DE results [36]. At least half of differential expression based studies incorporate a fold change cut-off typically between 2-3, however, informative RNAs and expressed transcripts have been shown to have a fold change less than 2 [37], and genes with low fold change have been demonstrated to influence biological effects in signalling cascades and pathways [36]. In addition, gene expression is heavily influenced by tissue, and as this study performs meta-analysis across multiple inter-related tissues within larger brain compartments, we do not employ an arbitrary fold change cut-off to determine if a gene is differentially expressed, however, we do require the gene to be consistently expressed across these tissues. if a gene was significantly DE according to the meta-analysis (FDR adjusted meta p-value ≤ 0.05), but at least one contributing dataset (according to AW.OC weights) had directional logFC discrepancy (i.e. up-regulated in one dataset and down-regulated in another dataset), the gene was deemed to be discordant and was excluded from further analysis. This ensured we only captured robust, and consistently reproducible expression signatures.

### Generation of disease-specific meta-analysis expression profiles

Meta-analysis was performed across all AD datasets, followed by a separate meta-analysis across the non-AD disorder datasets. Using these meta-analysis results we generated three expression profiles; (1) “AD expression profile”, (2) “AD-specific expression profile” and (3) “common neurological disorder expression profile”.

The first expression profile, “AD expression profile”, is a direct result of the meta-analysis performed on AD studies, which represents the changes typically observed from an AD and cognitively healthy control study design. The second expression profile, deemed as the “AD-specific expression profile”, is produced by subtracting significantly DEG’s found in the non-AD meta-analysis results from the “AD expression profile”. This profile represents transcriptomic changes specifically observed in AD and not in any other neurodegenerative or mental health disorder used in this study. The third expression profile, deemed as the “common neurological disorder expression profile”, represents genes which are significantly DE in all disorders used in this study, including AD.

### Replication of significant microarray genes in RNA-seq data

The genes significantly DE and deemed to be of biological significance in this study were queried in the curated web-based database Agora (data version 9, accessible at https://agora.ampadportal.org), which provides expression change of genes in AD based on RNA-seq of 2100 human brain samples.

### Functional and gene set enrichment analysis

Gene set enrichment analysis (GSEA), and Gene Ontology (GO) analysis was conducted using an Over-Representation Analysis (ORA) implemented through the ConsensusPathDB web platform (version 32) [38] in May 2017. ConsensusPathDB incorporates numerous well-known biological pathway databases including BioCarta, KEGG, Reactome and Wikipathways. The platform performs a hypergeometric test while integrating a background gene list, which in this case is a list of all the genes that pass quality control in this study, compiles results from each database and corrects for multiple testing using the false discovery rate (FDR) [38]. A minimum overlap of the query signature and database was set to 2, and a result was deemed significant if the q-value was ≤ 0.05.

### Network analysis

Protein-protein interaction (PPI) networks were created by uploading the meta-analysis DEG lists (referred to as seeds in network analysis) along with their meta logFC expression values to NetworkAnalyst’s web-based platform http://www.networkanalyst.ca/faces/home.xhtml in June 2017. The “Zero-order Network” option was incorporated to allow only seed proteins directly interacting with each other, preventing the well-known “Hairball effect” and allowing for better visualisation and interpretation [39]. Sub-modules with a p-value ≤ 0.05 (based on the “InfoMap” algorithm [40]) were considered significant key hubs, and the gene with the most connections within this hub was regarded as the key hub gene.

## RESULTS

### The AD microarray datasets

We Identified and acquired nine publicly available AD studies from ArrayExpress and AMP-AD, of which seven studies contained samples extracted from differing regions of the brain. The basic characteristics of each study and dataset are provided in Tables

Table 1. Separating the nine studies by brain regions resulted in 46 datasets. Here a “dataset” is defined by brain region and study origin. For example, ArrayExpress study E-GEOD-36980 consists of diseased and healthy samples extracted from three different tissues (temporal cortex, hippocampus and frontal cortex). All samples originating from the same tissue were classified as one dataset; therefore, study E-GEOD-36980 generated three datasets, representing the three different tissues. The 46 AD datasets contained both AD samples and healthy controls, and were assayed using seven different expression chips over two different microarray platforms (Affymetrix and Illumina) and consisted of a total 2718 samples before QC. Briefly, the MetaOmics analysis identified study syn4552659 as an outlier and was therefore removed from further analysis (see supplementary text 1), resulting in 1501 samples (746 AD, 755 controls) in the remaining 22 datasets after QC

**Table 1:**
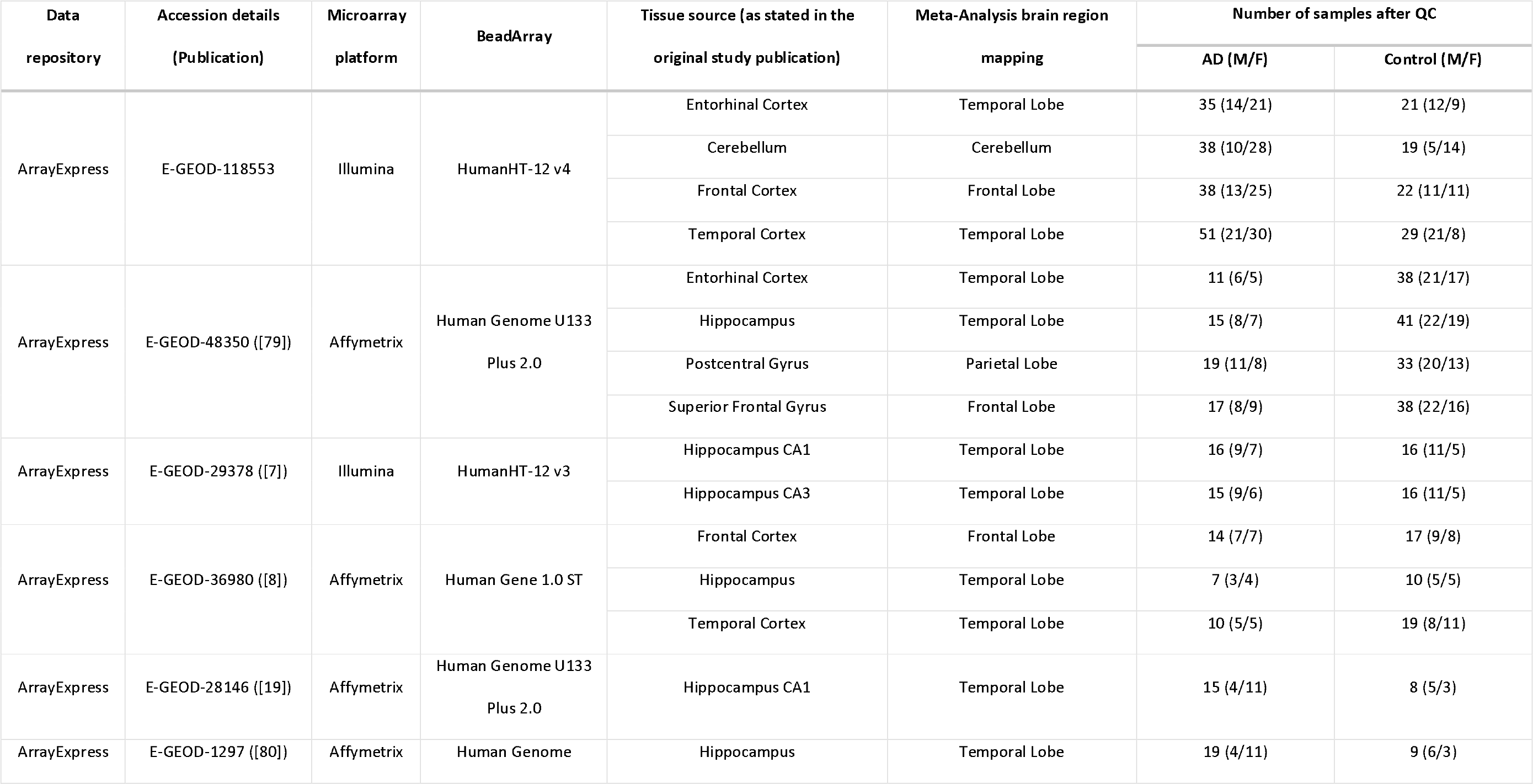
Characteristics of individual AD studies processed in this meta-analysis

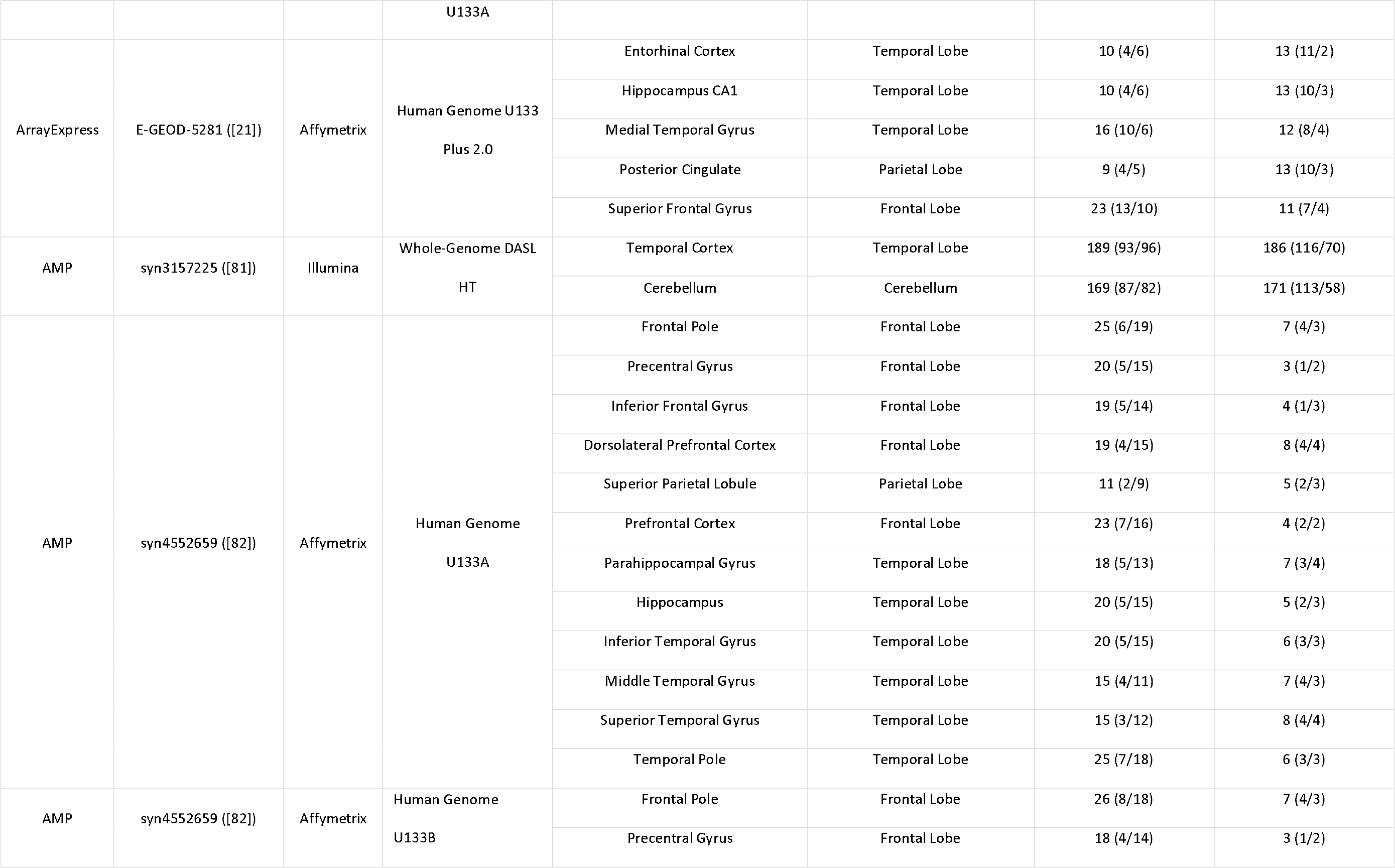

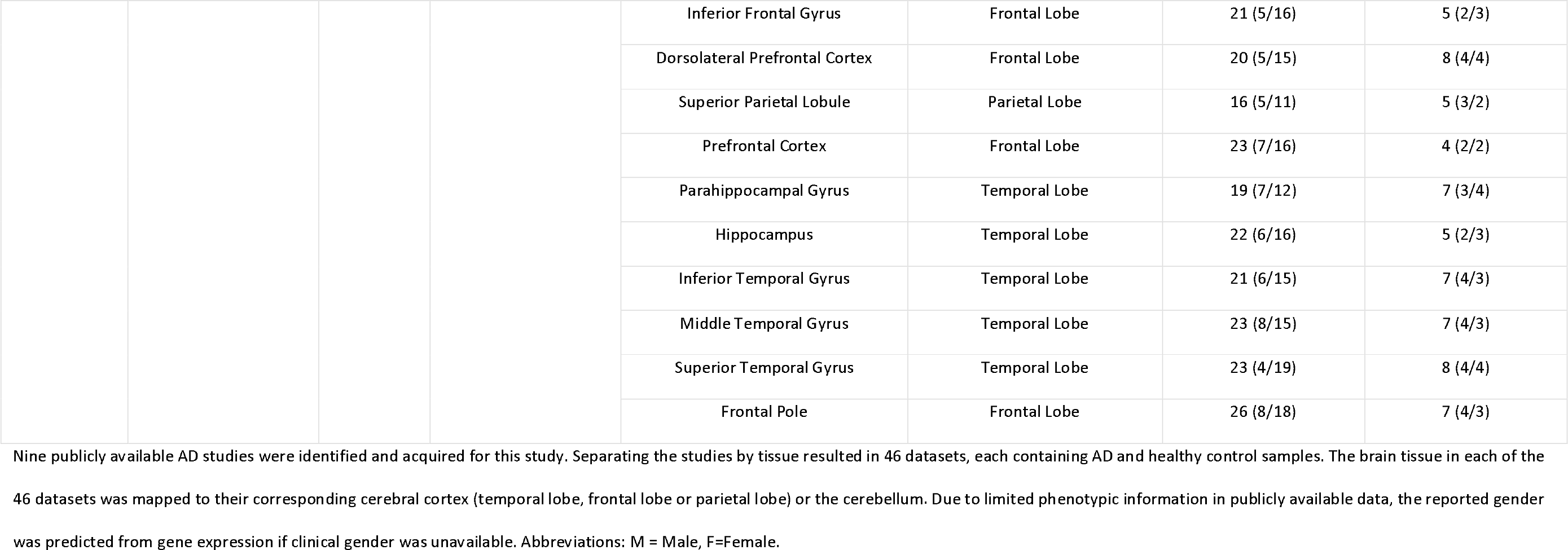

### Summary of the AD meta-analysis DEG counts

The AD meta-analysis was performed on the 22 AD datasets and independently identified differentially expressed genes within the TL, FL, PL and CB brain regions. A summary of the number of datasets in each brain region and the number of significant DEG’s identified is provided in Table 2**Error! Reference source not found.**. The complete DE results are provided in Supplementary Table 1.

**Table 2:** Summary of AD study meta-analysis DEG’s

| Brain region | Number of datasets | Number of samples<br>(case/control) | AW.OC Significant DEGs<br>(FDR adjusted $p \leq 0.05$) |
| --- | --- | --- | --- |
| Temporal lobe | 14 | 850 (419/431) | 323 |
| Frontal lobe | 4 | 180 (92/88) | 460 |
| Parietal lobe | 2 | 74 (28/46) | 1736 |
| Cerebellum | 2 | 397 (207/190) | 867 |

Twenty-two AD datasets containing a total of 1501 samples remained in this study after QC. The case/control numbers represent the total number of AD/healthy controls subjects across all datasets within a particular brain region. The number of significant genes was identified through a combining p-value method known as Adaptively Weighted with One-sided Correction (AW.OC).

### The non-AD disorder microarray datasets

Nine non-AD studies were identified and acquired, of which four studies consisted of samples generated from multiple disorders and brain regions. Separating the studies by disease and tissue equated to 21 datasets consisting of 8 SCZ, 6 BD, 4 HD, 2 MDD and 1 PD dataset with a total of 1166 samples after QC. The demographics of the non-AD datasets is provided in Table 3

**Table 3:**
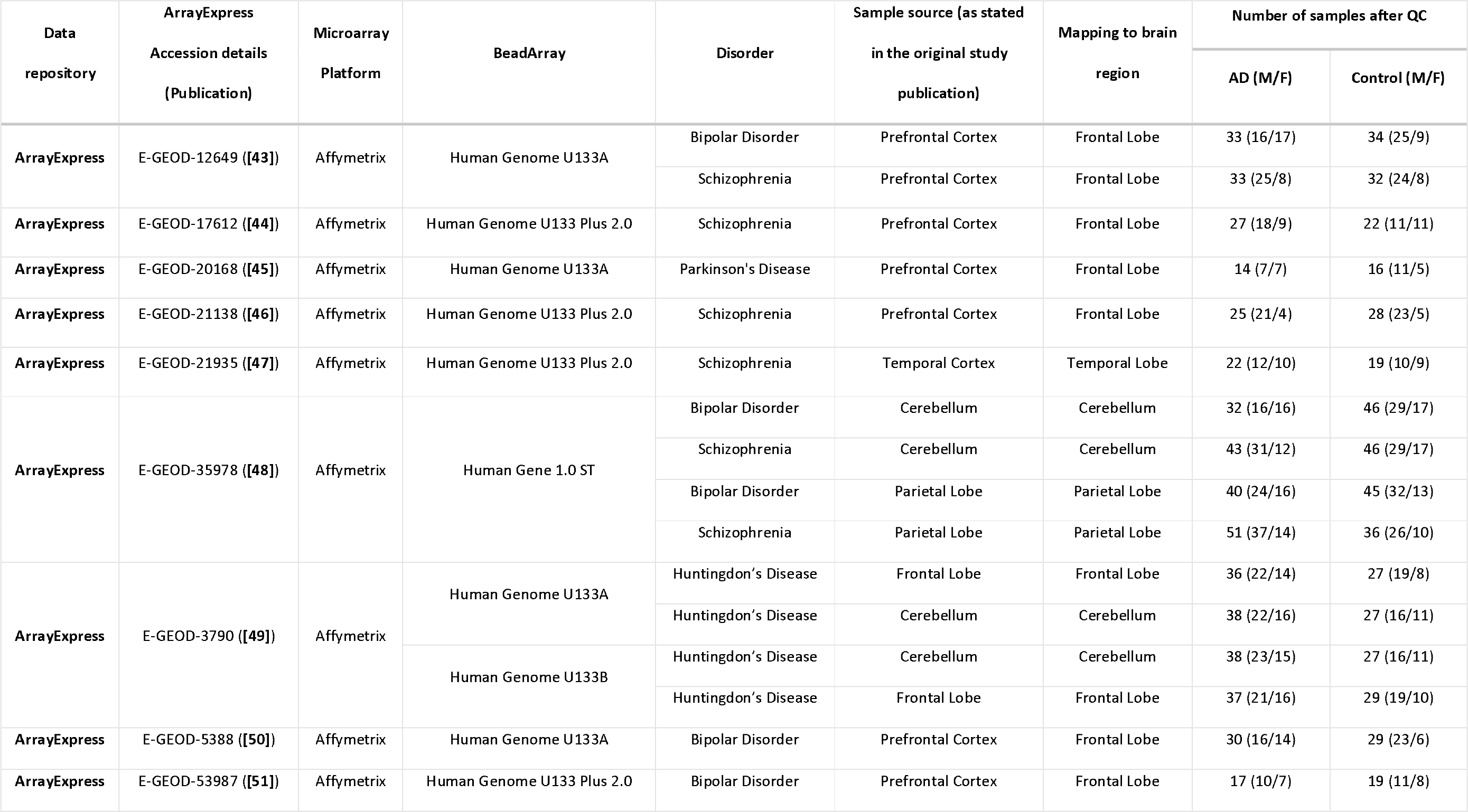
Characteristics of individual non-AD studies included in this meta-analysis

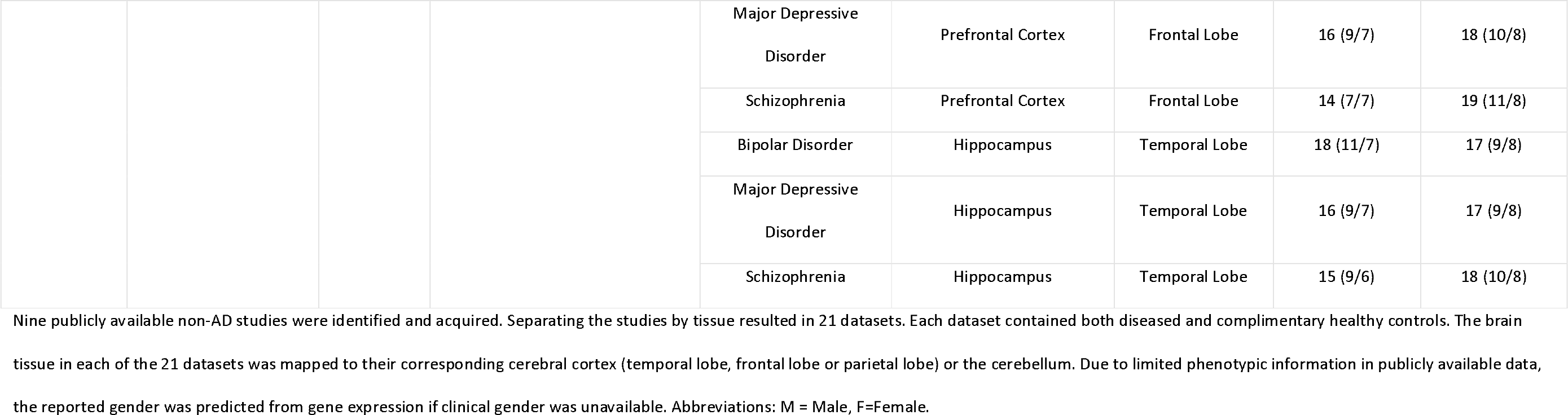

### Summary of non-AD brain disorder meta-analyses DEG counts

A second meta-analysis was performed on all non-AD disorders, and similarly to the AD meta-analysis, datasets were grouped into the TL, FL, PL and CB brain regions. An overview of the non-AD meta-analysis results are provided in Table 4, and a complete list of DEG’s is provided in Supplementary Table 2. SCZ and BD were the only disorders with expression data available across all four brain regions, and the frontal lobe brain region was the only region with expression data available from all non-AD disorders identified in this study.

**Table 4:** Summary of non-AD study meta-analysis DEG’s

| Brain region | Number of<br>BD datasets<br>(case/control) | Number of<br>Schizophrenia<br>datasets<br>(case/control) | Number of HD<br>datasets<br>(case/control) | Number of<br>MDD datasets<br>(case/control) | Number of PD<br>datasets<br>(case/control) | Total number of<br>datasets<br>(case/control) | AW.OC Significant<br>DEGs (FDR adjusted<br>p ≤ 0.05) |
| --- | --- | --- | --- | --- | --- | --- | --- |
| Temporal lobe | 1 (18/17) | 2 (37/37) | 0 | 1 (16/17) | 0 | 4 (71/71) | 51 |
| Frontal lobe | 3 (80/82) | 4 (99/101) | 2 (73/56) | 1 (16/18) | 1 (14/16) | 11 (282/273) | 149 |
| Parietal lobe | 1 (40/45) | 1 (51/36) | 0 | 0 | 0 | 2 (91/81) | 2611 |
| Cerebellum | 1 (32/46) | 1 (43/46) | 2 (76/54) | 0 | 0 | 4 (151/146) | 177 |

The table illustrates the non-AD dataset and sample distribution across the four brain regions. Disease abbreviations are as follows:
BD=Bipolar Disease, HD= Huntington's Disease, MDD=Major Depressive Disorder and PD=Parkinson's Disease. The case/control numbers represent the total number of diseased and healthy control subjects within a disease group and brain region. For instance, "3 (80/82)" for BD datasets in the Frontal lobe region indicates three BD datasets with a combined total of 80 BD and 82 complimentary healthy control subjects. The number of significant DEG's was identified through a combining p-value method known as Adaptively Weighted with One-sided Correction (AW.OC).

### The meta-analysis expression profiles

As described in the methods, three primary expression signatures were derived from the meta-analyses for each of the four brain regions: - 1) “AD expression profile”, 2) “AD-specific expression profile” and 3) “common neurological disorder expression profile”. The numbers of significant DEG’s in each of the three expression signatures are provided in Table 5.

**Table 5:**
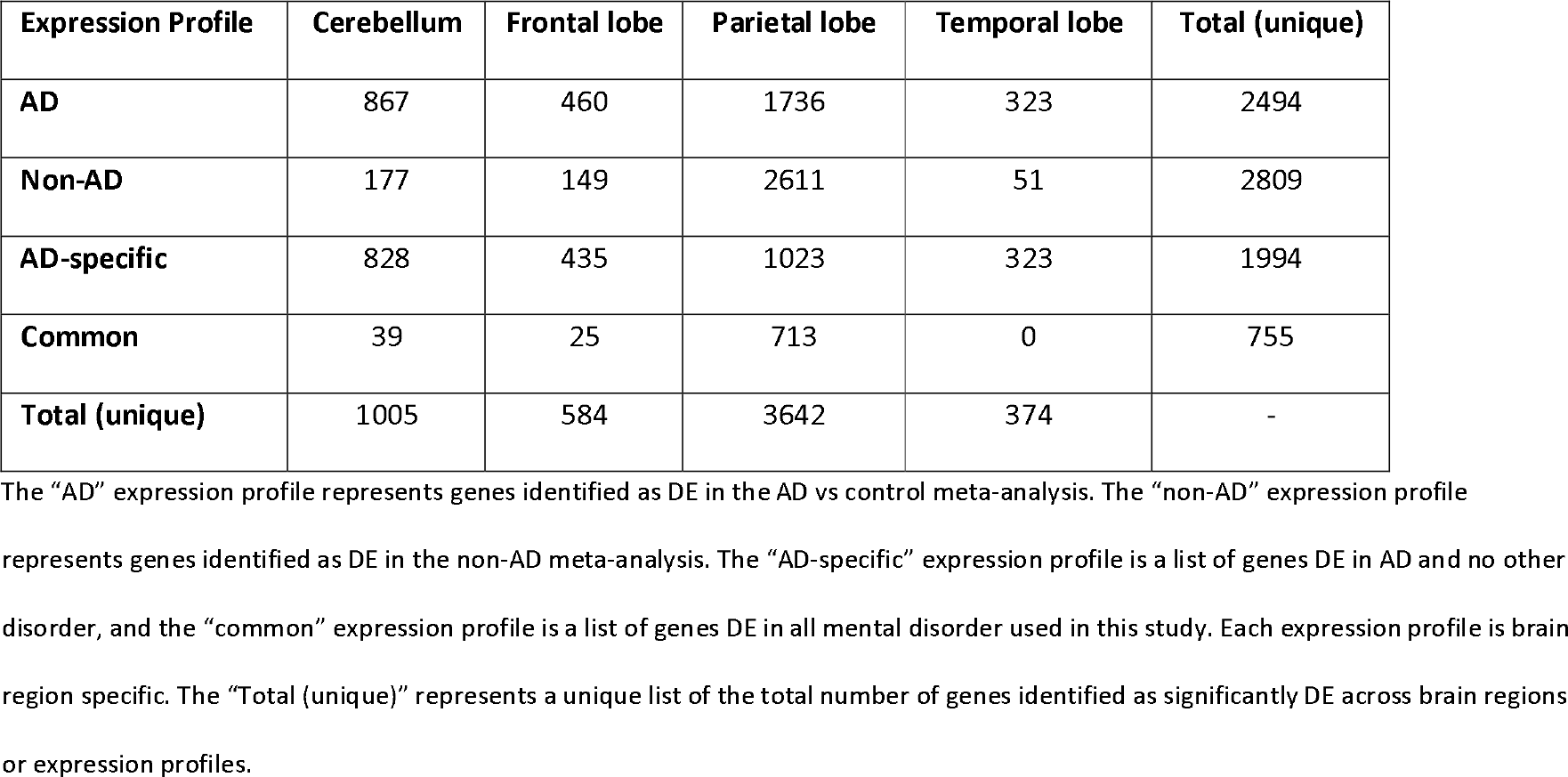
Summary of DEGs in each expression signature and brain region

The DEG’s from the “AD expression profile” in the TL brain region were not significantly DE in any other disorder included in this study. Hence, the “AD expression profile” and the “AD-specific expression profile” contained the same 323 genes for the TL brain region. The “AD-specific expression profile” for all four brain regions is provided in Supplementary Table 3.

The “common neurological disorder expression profile” within the four brain regions consisted of very little or no DEG’s (except for the parietal lobe); hence, the downstream analysis did not yield any statistically significant results of biological relevance. We find little robust evidence of shared biology based on this data analysis and therefore, exclude all results generated from the “common neurological disorder expression profile” from this paper; however, we provide the complete list of significantly DEG’s within this profile in Supplementary Table 4.

### Common differentially expressed genes across multiple brain regions in AD

AD is known to affect all brain regions through the course of the disease, although not to the same degree, similar transcriptomic changes across all brain regions were deemed disease-specific, while perturbations in a single brain region were considered to be tissue-specific. We were particularly interested in disease-specific transcriptomic changes and therefore decided to focus on genes that were found to be consistently DE across multiple brain regions.

Meta-analysis of the AD datasets identified a total of 2495 unique genes as significantly DE. The distribution of these genes across the four brain regions is shown in Figure 1. Figure 1 orty-two genes were found to be perturbed across all four brain regions and can be grouped into three sets (Figure 2). The first group (Gene set 1) are expressed consistently in the same direction across all four brain regions and can be regarded as disease-specific. The second group (Gene set 2) are expressed in the same direction in the TL, FL and PL, but expression is reversed in the CB brain region, a region suggested to be spared from AD pathology [4] [5]. This expression pattern suggests these genes may be involved in AD pathology. Finally, the third group (Gene set 3) are inconsistently expressed across the four brain regions are most likely tissue-specific or even false-positives.

**Figure 1:**
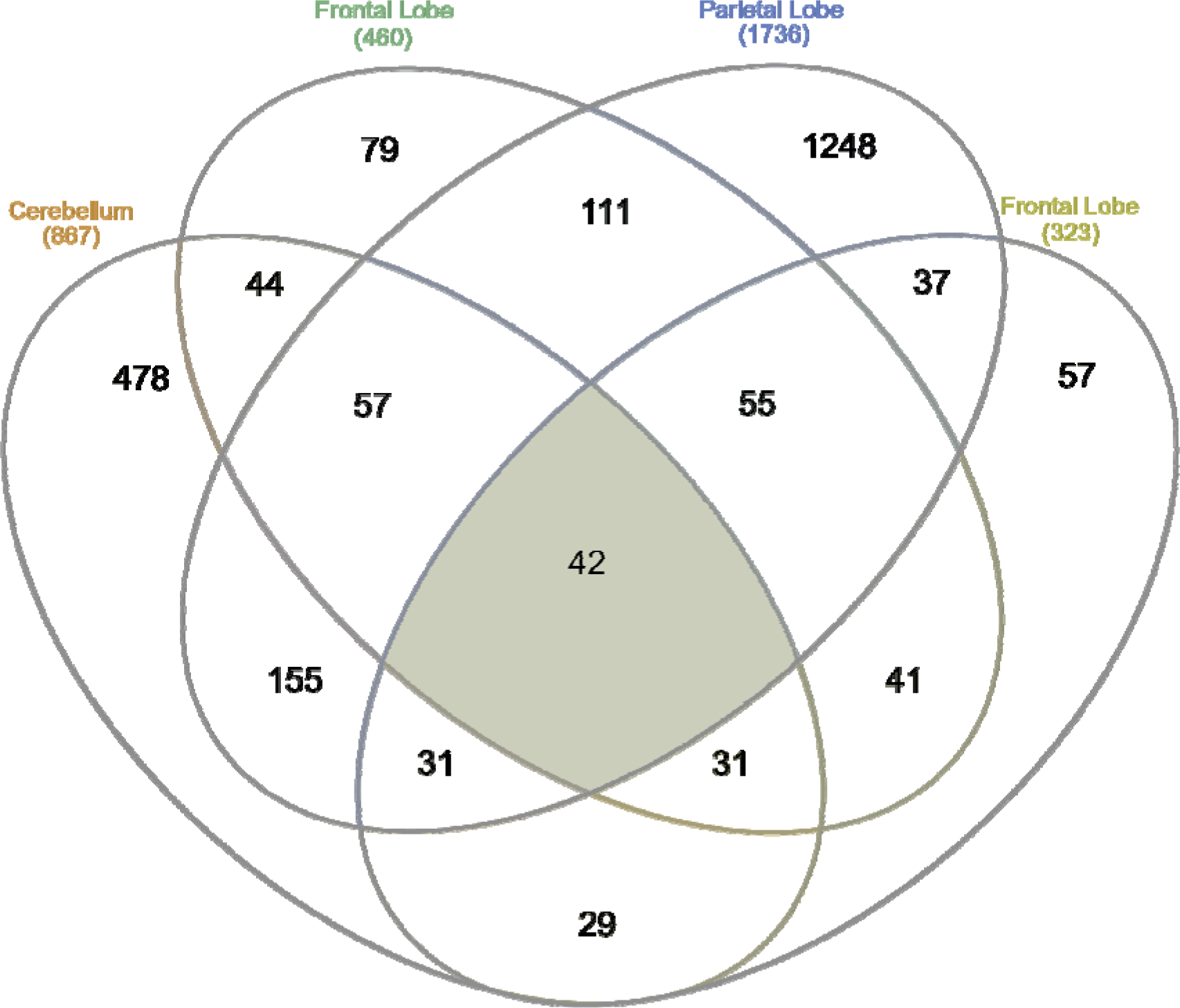
Overlap of DEG’s in the AD expression profile across brain regions. Forty-two genes were observed to be significantly differentially expressed across all four AD brain regions.

**Figure 2:**
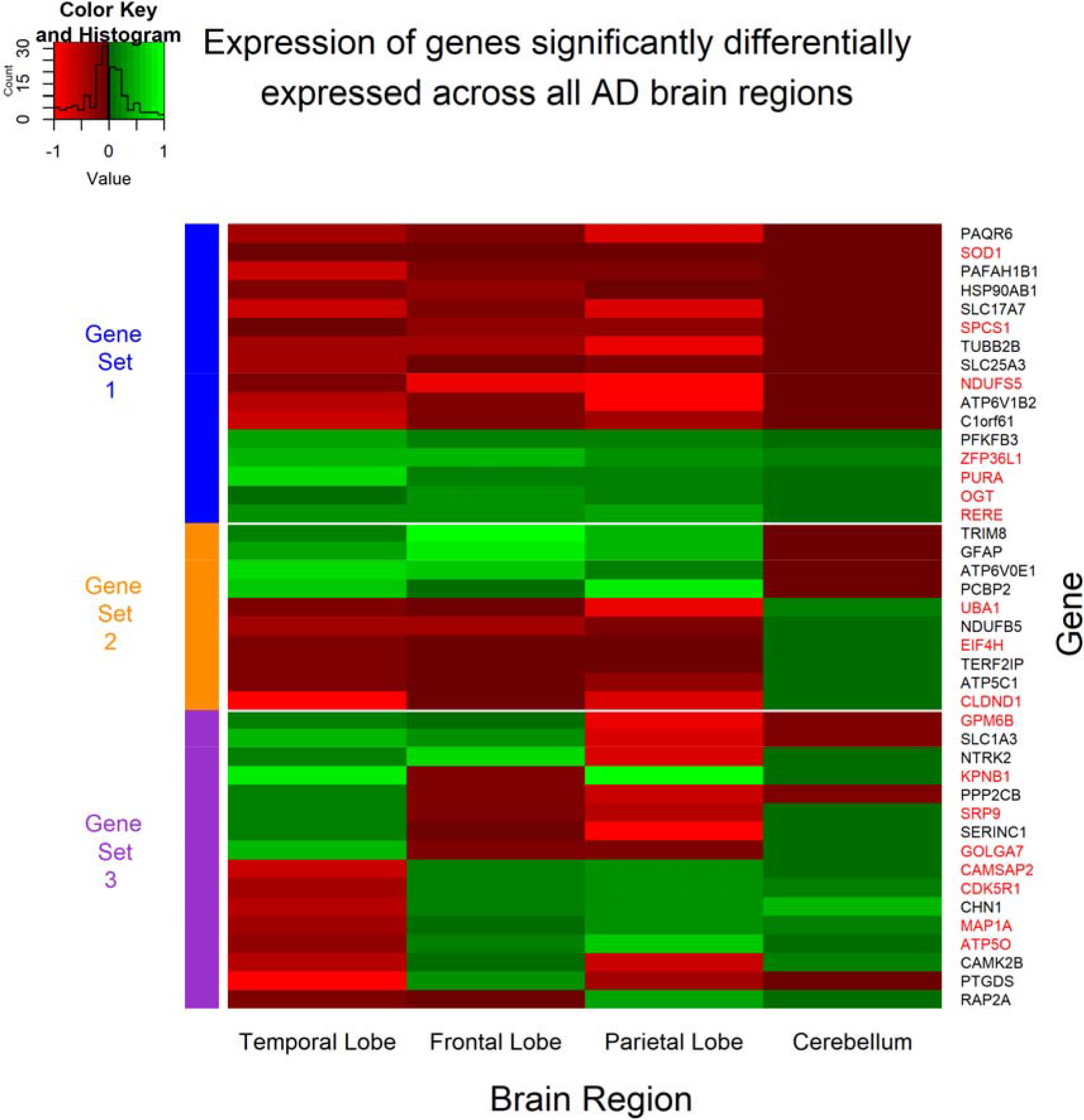
Expression pattern of genes significantly differentially expressed across all four AD brain regions. The expression values for each gene was obtained from the meta-summary calculations. Red cells represent down-regulated genes, and green cells represent up-regulated genes. Forty-two genes were observed to be significantly perturbed across all four AD brain regions and can be grouped into three “sets”. Gene set 1 represents genes which are perturbed consistently in the same direction across all AD brain regions and can be considered disease-specific. Gene set 2 represents genes consistent in expression in the TL, FL and PL brain regions, but reversed in the CB brain region; a region often referred to be free from AD pathology. Finally, Gene set 3 represents genes which are significant DE across all four brain regions, however, directional change is not consistent across the brain regions and may represent tissue-specific genes or even false positive. The gene names highlighted in red are genes perturbed in AD and not in any other disorder used in this study and are deemed “AD-specific”.

From the forty-two genes significantly differentially expressed across all brain regions, seven genes were DE in the same direction and belong to the “AD-specific expression profile”, that is, these seven genes (down-regulated **NDUFS5, SOD1, SPCS1** and up-regulated **OGT, PURA, RERE, ZFP36L1**) were consistently perturbed in all AD brain regions and not in any other brain region of any other disorder used in this study and can be considered unique to AD brains. The expression of these seven genes across AD brains is shown in Figure 3.

**Figure 3:**
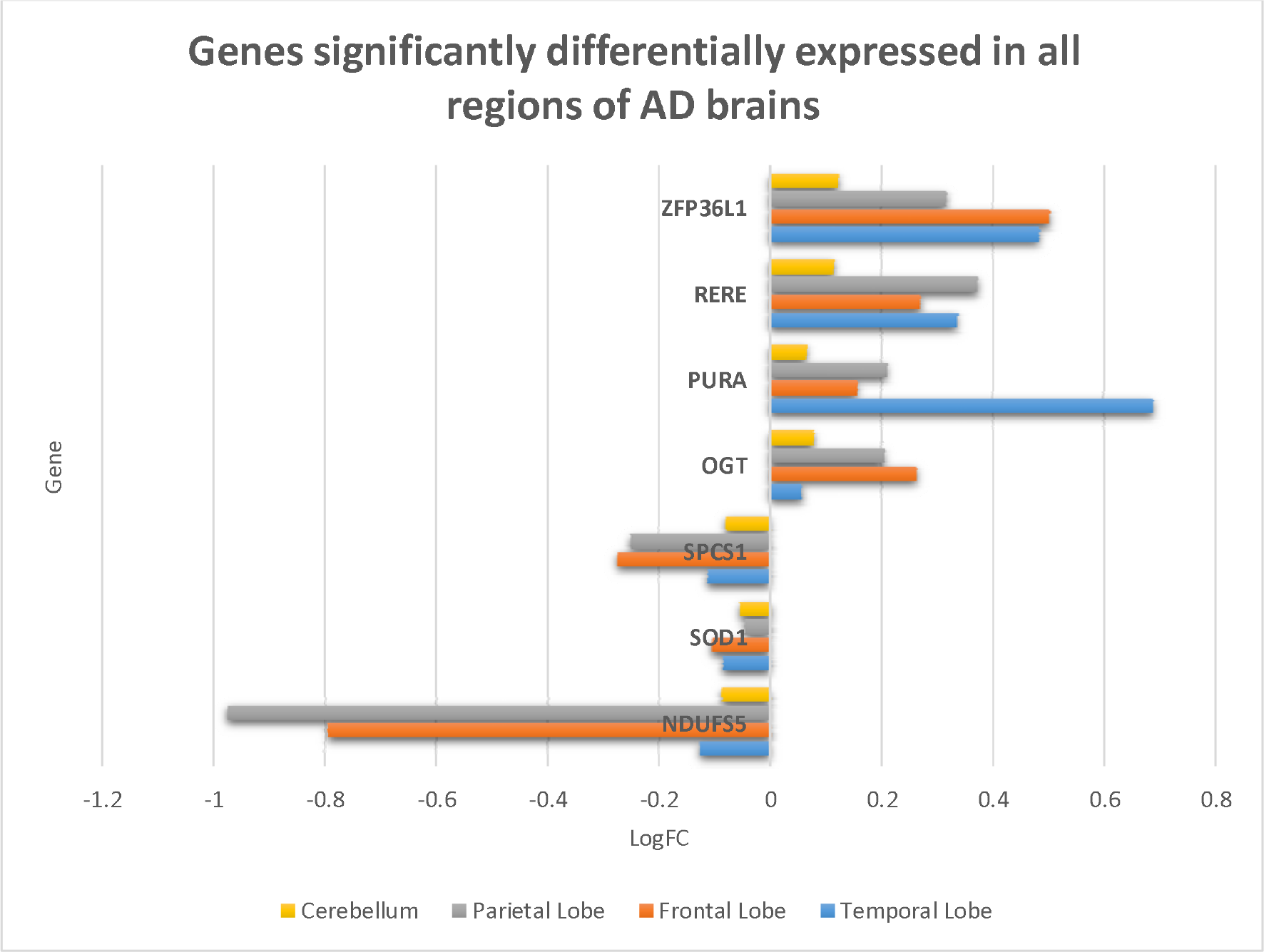
Seven genes consistently significantly differentially expressed in the same direction in all regions of AD brains but not in Schizophrenia, Bipolar Disorder, Huntington’s disease, Major Depressive Disorder or Parkinson’s disease brains. These seven genes can be assumed to be unique to AD brains, and may play an important role in disease mechanisms.

### Differentially expressed genes in brain regions affected by AD histopathology

In AD, the TL, FL and PL are known to be affected by both plaques and tangles, while the CB brain region is rarely reported to be affected. In addition to identifying genes DE across all brain regions and reversed in the CB brain region, we were also interested in genes perturbed in the TL, FL and PL and not the CB. These genes may also play a role in general AD histopathology and could be new therapeutic targets in preventing or curing AD.

Fifty-five genes were found to be significantly DE in TL, FL and PL but not the CB, of which sixteen were expressed in the same direction and were not DE in the other brain disorders used in this study. Ten of these genes (**ALDOA, GABBR1, TUBA1A, GAPDH, DNM3, KLC1, COX6C, ACTG1, CLTA, SLC25A5**) were consistently down-regulated, and six genes (**PRNP, FDFT1, RHOQ, B2M, SPP1, WAC**) were consistently up-regulated in AD.

Furthermore, from the forty-two genes identified as significantly DE across all four AD brain regions, ten genes were in consensus in their expression across the TL, FL and PL brain region but expression is reversed in the cerebellum. Only 3 of these genes (**UBA1, EIF4H** and **CLDND1**) belong to the “AD-specific expression profile”, and all three genes were significantly down-regulated in the TL, FL and PL, but significantly up-regulated in the CB brain region (see Gene set 2 in Figure 2).

### Microarray gene expression profiling in RNA-seq data

The 7 genes (**NDUFS5, SOD1, SPCS1, OGT, PURA, RERE, ZFP36L1**) consistently expressed across all brain regions and the 19 genes (**ALDOA, GABBR1, TUBA1A, GAPDH, DNM3, KLC1, COX6C, ACTG1, CLTA, SLC25A5, PRNP, FDFT1, RHOQ, B2M, SPP1, WAC, UBA1, EIF4H, CLDND1**) consistently expressed in the TL, FL and PL and not in the CB or reversed in the CB, were queried in the web-based platform Agora to compare RNA-seq based expression profiling. The results are provided in Table 6. Agora failed to provide expression profiling for 17/26 genes, however, from the data available, the genes observed to be consistently expressed across all brain regions based on microarray data are relatively mirrored in RNA-seq data, specifically genes **SPCS1, PURA** and **ZFP36L1**.

**Table 6:**
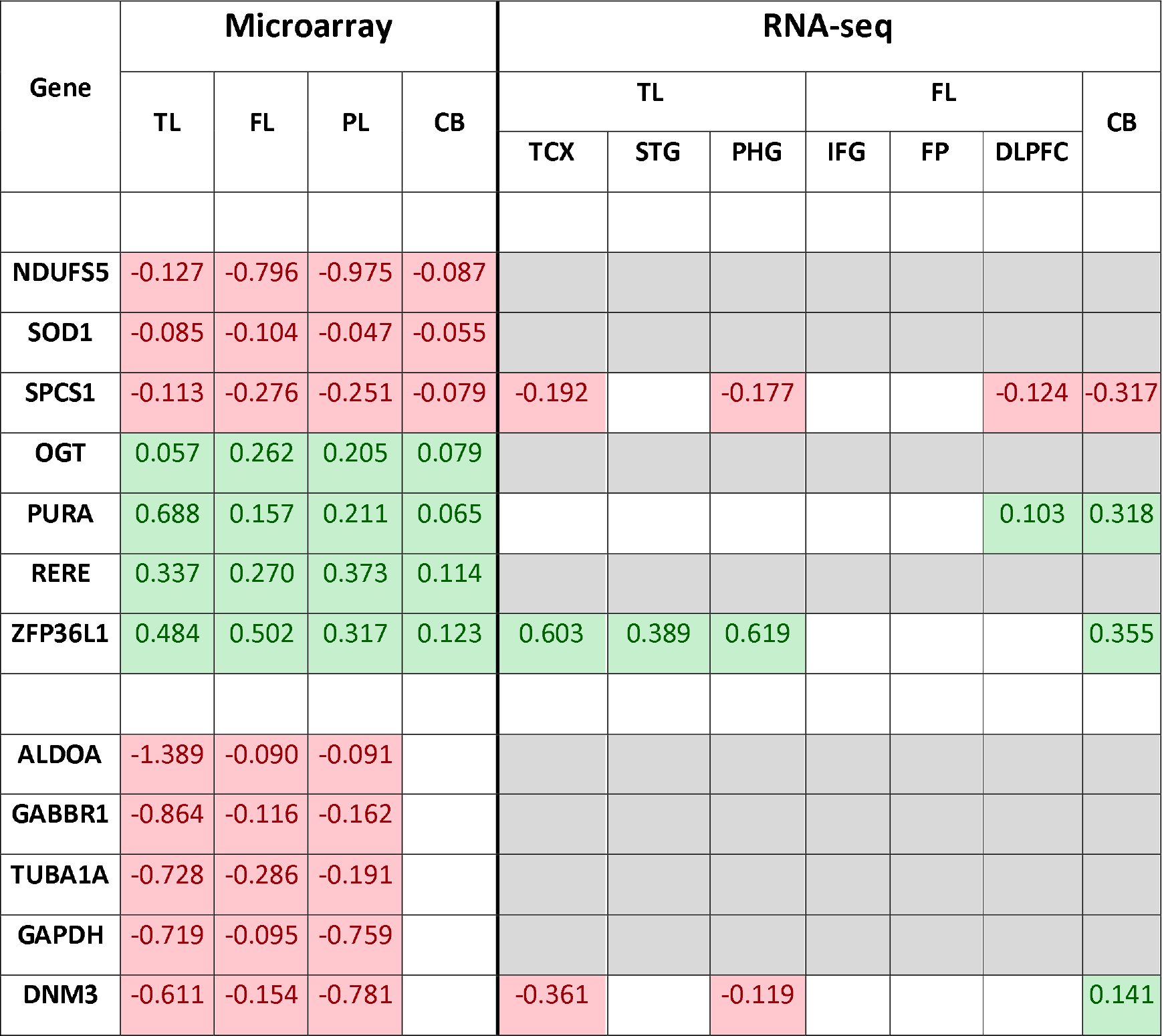
Microarray gene expression compared to RNA-seq gene expression

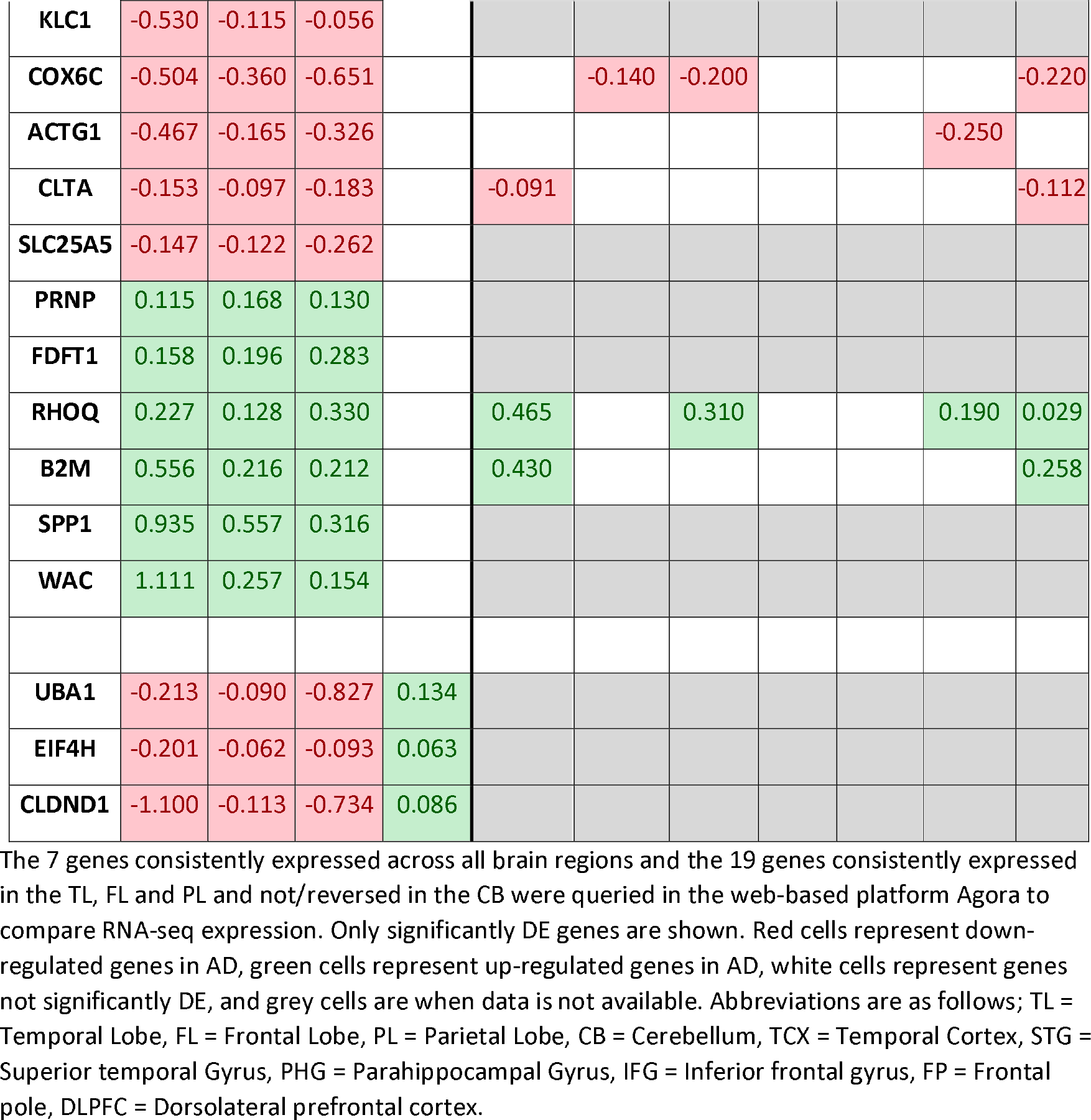

RNA-seq data was available for only 6/19 genes (DNM3, COX6C, ACTG1, CLTA, RHOQ and B2M) expressed in brain regions affected by hallmark AD pathology (TL, FL and PL), and were all relatively consistent in significance and directional change across AD brain regions, including the CB, a characteristic undesired by genes which may be associated with hallmark AD pathology.

### “AD Expression Profile” functional gene set enrichment and GO analysis

Gene set enrichment analysis of the “AD expression profile” identified 205, 197, 98, and 45 biological pathways significantly enriched in the TL, FL, PL and CB brain regions respectively (Supplementary Table 5). There were ten pathways significantly enriched in all four brain regions, of which eight are involved in the “**metabolism of protein**” (specifically the translation process, the most significant being in CB brain region with a q-value=1.11e-7), one involved in “**adenosine ribonucleotides de novo biosynthesis**” (TL q-value = 0.007, FL q-value = 7.56e-5, PL q-value = 0.04, CB q-value = 0.03) and one involved in the “**digestive system**” (TL q-value = 0.02, FL q-value = 0.02, PL q-value = 0.01, CB p-value = 0.02).

When excluding the CB brain region, 42 pathways were significantly enriched in the remaining three brain regions, of which five pathways obtained an FDR adjusted significance p-value of ≤ 0.01. The five pathways are “**Alzheimer’s disease**” (TL q-value = 6.53e-4, FL q-value = 0.02, PL q-value = 0.01), “**Electron Transport Chain**” (TL q-value = 0.006, FL q-value = 2.95e-5, PL q-value = 3.69e-5), “**Oxidative phosphorylation**” (TL q-value =1.77e-4, FL q-value = 4.99e-8, PL q-value = 4.18e-05), “**Parkinson’s disease**” (TL q-value =8.57e-4, FL q-value = 1.59e-6, PL q-value = 1.77e-6) and “**Synaptic vesicle cycle**” (TL q-value = 5.19e-4, FL q-value = 3.82e-7, PL q-value = 2.03e-4).

The biological GO analysis identified 384, 417, 216, and 72 biological components as significantly enriched in the TL, FL, PL and CB brain region respectively (Supplementary Table 6). There were 36 pathways significantly enriched across all four brain regions at a p-value threshold of ≤ 0.05 and nine at an FDR adjusted significant p-value threshold of ≤ 0.01. These nine processes are “**cellular component biogenesis**” (TL q-value =1.38e-4, FL q-value = 0.002, PL q-value = 5.86e-4, CB q-value = 0.006), “**cellular component organization**” (TL q-value = 1.96e-8, FL q-value = 1.04e-8, PL q-value = 3.35e-5, CB q-value = 0.004), “**interspecies interaction between organisms**” (TL q-value = 1.13e-41.85e-4, FL q-value = 8.73e-5, PL q-value = 5.59e-5, CB q-value = 0.002), “**multi-organism cellular process**” (TL q-value =, FL q-value = 4.72e-5, PL q-value = 8.04e-5, CB q-value = 0.002), “**nervous system development**” (TL q-value = 1.64e-7, FL q-value = 5.90e-14, PL q-value = 3.82e-8, CB q-value = 0.01), “**organonitrogen compound metabolic process**” (TL q-value = 0.002, FL q-value = 1.56e-5, PL q-value = 1.02e-5, CB q-value = 0.002), “**symbiosis, encompassing, mutualism through parasitism**” (TL q-value = 4.04e-4, FL q-value = 1.92e-4, PL q-value = 3.18e-4, CB q-value = 0.004), “**translational initiation**” (TL q-value = 0.007, FL q-value = 0.006, PL q-value = 2.41e-4, CB q-value = 5.24e-6), and “**viral process**” (TL q-value = 2.82e-4, FL q-value = 1.17e-4, PL q-value = 3.18e-4, CB q-value = 0.002). Excluding the CB brain region resulted in 84 common biological components being significantly enriched across the remaining three brain regions.

### “AD-Specific Expression Profile” functional gene set enrichment and GO analysis

Analysis of the “AD-specific expression profile” identified 205, 196, 40 and 42 pathways as significantly enriched in the TL, FL, PL and CB brain region respectively in the GSEA analysis (Supplementary Table 7). The analysis identified six significantly enriched pathways across all four brain regions, and all are involved in “**metabolism of protein**” (specifically the translation process, with the most significant pathway being in the PL brain region with a q-value = 8.92e-7). The same six pathways were identified when the CB region was excluded.

The GO analysis identified 384, 344, 36 and 72 significantly enriched biological components for the TL, FL, PL and CB brain region respectively. Only four common biological components were significantly enriched across all four brain regions, and all are indicative of interspecies interactions including viral. Excluding the CB identifies only “**neural nucleus development**” (TL q-value = 5.35e-5, FL q-value = 0.007, PL q-value = 0.003) as an additional component being enriched. The complete biological GO analysis results are provided in Supplementary Table 8.

### Network analysis hub gene identification

PPI networks were generated for each expression profile and in each of the four brain regions (TL, FL, PL and CB) to identify genes whose protein product interacts with other protein products from the same expression profile. Genes with more interactions than expected are referred to as hub genes and may be of biological significance.

### Temporal lobe hub genes

PPI network analysis was performed on the expression profiles of TL brain region to identify key hub genes. The “AD expression profile” and the “AD-specific expression profile” both consisted of the same 323 DEG’s which represented 282 seed proteins with 716 edges (interactions between proteins). Two significant key hub genes were identified; the down-regulated Polyubiquitin-C (**UBC**, p-value = 1.57e-30) and the up-regulated Small Ubiquitin-related Modifier 2 (**SUMO2**, p-value = 3.7e-4).

### Frontal Lobe hub genes

The FL “AD expression profile” consisted of 460 DEG which represented 272 seed proteins and 620 edges. Two significant key hub genes were identified; up-regulated Amyloid Precursor Protein (**APP**, p-value = 1.98e-08) and down-regulated Heat Shock Protein 90-alpha (**HSP90AA1**, p-value = 0.003). Using the “AD-specific expression profile” identified the same two key hub genes, with **APP** reaching a significant p-value of 2.11e-09.

### Parietal Lobe hub genes

The PL “AD expression profile” consisted of 1736 DEG which represented 1437 seed proteins and 5720 edges. Similar to the TL and FL, two significant key hub genes were identified; down-regulated Cullin-3 (**CUL3**, p-value = 1.84e-10) and down-regulated **UBC** (p-value = 1.84e-10). Using the “AD-specific expression profile” (1023 DEGs, 810 seed proteins and 2351 edges) identified **UBC** as the only key hub gene, with a more significant p-value of 1.84e-10. The **CUL3** gene is no longer a significant key hub gene in the network.

### Cerebellum hub genes

The CB “AD expression profile” consisted of 867 DEG’s which represented 548 seed proteins and 1419 edges. Four significant key hub genes were identified; up-regulated **APP** (p-value =4.24e-26), down-regulated Ribosomal Protein 2 (**RPS2**, p-value = 4.24e-26), down-regulated **SUMO2** (p-value = 4e-05), and up-regulated Glycyl-TRNA Synthetase (**GARS**, p-value = 0.0207). Using the “AD-specific expression profile” for the same brain region identified **APP** (p-value = 3.44e-26), **RPS2** (p value= 6.61e-06), and **SUMO2** (p-value = 3.78e-06) as the key hub genes only. The **GARS** gene is no longer a key hub gene in the network.

## DISCUSSION

In this study, we acquired eighteen publicly available microarray gene expression studies covering six neurological and mental health disorders; AD, BD, HD, MDD, PD and SCZ. Data was generated on seven different expression BeadArrays and across two different microarray technologies (Affymetrix and Illumina). The eighteen studies consisted of 3984 samples extracted from 22 unique brain regions which equated to 67 unique datasets when separating by disorder and tissue. However, due to study and sample outlier analysis, only 43 datasets (22 AD, 6 BD, 4 HD, 2 MDD, 1 PD and 8 SCZ) totalling 2,667 samples passed QC. We grouped the AD datasets by tissue, into the TL, FL, PL and CB brain regions to perform the largest microarray AD meta-analysis known to date to our knowledge, which identified 323, 460, 1736 and 867 significant DEG’s respectively. Furthermore, we incorporated transcriptomic information from other neurological and mental health disorders to subset the initial findings to 323, 435, 1023, and 828 significant DEG’s that were specifically perturbed in the TL, FL, PL and CB brain regions respectively of AD subjects.

### Genes specifically perturbed across AD brain regions

Seven genes (down-regulated **NDUFS5, SOD1, SPCS1** and up-regulated **OGT, PURA, RERE, ZFP36L1**) were DE in AD brains and not DE in the other disorders used in this study. We deemed these seven protein-coding genes as “AD-specific”. The expression patterns of three genes (**SPCS1, PURA** and **ZFP36L1**) were relatively mirrored in RNA-seq data, however, it is important to note the RNA-seq data does not contain expression profiling for the PL region and it also contains three specific brain regions within the TL (temporal cortex, superior temporal gyrus, Parahippocampal gyrus) and FL (Inferior frontal gyrus, frontal pole and dorsolateral prefrontal cortex). Nevertheless, SPCS1 gene was observed to be consistently down-regulated across all hierarchical AD brain regions available in both the microarray and RNA-seq data. In addition, based on a network of genomics and epigenomic elements in the region of this genes, in combination with phenotypes, the AMP-AD consortia have nominated **SPCS1** as a druggable target for AD treatment, providing confidence the remaining genes identified in this study may also provide druggable targets.

Three of the “AD-specific” genes (**NDUFS5, SOD1** and **OGT**) have been previously associated with AD. Down-regulated NADH Dehydrogenase Ubiquinone Fe-S Protein 5 (**NDUFS5**) gene is part of the human mitochondrial respiratory chain complex; a process suggested to be disrupted in AD in multiple studies [38] [39]. A study investigating blood-based AD biomarkers identified 13 genes, including **NDUFS5**, which was capable of predicting AD with 66% accuracy (67% sensitivity and 75% specificity) in an independent cohort of 118 AD and 118 control subjects [43]. The perturbation in **NDUFS5** expression in the blood and brains of AD subjects suggests this gene may have potential as an AD biomarker and warrants further investigation.

Down-regulated Superoxide Dismutase 1 (**SOD1**) gene encodes for copper and zinc ion binding proteins which contribute to the destruction of free superoxide radicals in the body and is also involved in the function of motor neurons [provided by RefSeq, Jul 2008]. Mutations in this gene have been heavily implicated as causes of familial amyotrophic lateral sclerosis (**ALS**) [44] and have also been associated with AD risk [45]. A recent study discovered SOD1 deficiency in an amyloid precursor protein-overexpressing mouse model accelerated Aβ oligomerisation and also caused Tau phosphorylation [46]. They also stated **SOD1** isozymes were significantly decreased in human AD patients, and we can now confirm **SOD1** is significantly under-expressed at the mRNA level in human AD brains as well.

The up-regulated O-Linked N-Acetyl Glucosamine Transferase (**OGT**) gene encodes for a glycosyltransferase that links N-acetylglucosamine to serine and threonine residues (O-GlcNAc). O-GlcNAcylation is the post-translational modification of O-GlcNAc and occurs on both neuronal tau and APP. Increased brain O-GlcNAcylation has been observed to protect against tau and amyloid-β peptide toxicity [47]. A mouse study has demonstrated a deletion of the encoding **OGT** gene causes an increase in tau phosphorylation [48]. In this study, we observe a significant increase in **OGT** gene expression throughout human AD brains, including the cerebellum where tangles are rarely reported, suggesting **OGT** gene is most likely not solely responsible for the formation of tangles.

**OGT** and O-GlcNAcase (OGA) enzymes facilitate O-GlcNAc cycling, and levels of GlcNAc have also been observed to be increased in the parietal lobe of AD brains [49]. Appropriately, OGA inhibitors have been tested for treating AD with promising preliminary results [50], prompting further investigation into targeting **OGT** for AD treatment.

### Genes involved in AD histopathology

The CB brain region is known to be free from tau pathology and occasionally free from plaques. We exploited the CB brain region as a secondary control to identify sixteen genes (**ALDOA, GABBR1, TUBA1A, GAPDH, DNM3, KLC1, COX6C, ACTG1, CLTA, SLC25A5, PRNP, FDFT1, RHOQ, B2M, SPP1, WAC**) DE specifically in TL, FL and PL and not the CB brain region of AD subjects. RNA-seq data was available for 6 of these genes (**DNM3, COX6C, ACTG1, CLTA, RHOQ** and **B2M**) and all 6 genes failed to replicate expression patterns observed with microarray data. Nevertheless, DNM3 gene has been previously associated with AD pathology based on proteomic data DNM3 gene encodes a member of a family of guanosine triphosphate (GTP)-binding proteins that associate with microtubules and are involved in vesicular transport. A proteomic study identified a module of co-expressed proteins, which included DNM3, as negatively correlated with BRAAK staging [51]. Although DNM3 gene expression based on microarray and RNA-seq data are in disagreement in the CB brain region, a region used in this study to aid in determining whether a gene may be involved with AD pathology, an independent proteomic study demonstrated DNM3 might indeed be association with AD pathology. This suggests all 6 genes which failed replication in RNA-seq data may still be associated with AD pathology and require further confirmation.

An additional 9 genes (**GABBR1, GAPDH, PRPN, FDFT1, KLC1, TUBA1A, CLTA, COX6C, SLC25A5**), where expression profiling based on RNA-seq data was unavailable, have also been previously associated with AD, of which four genes (**GABBR1, GAPDH, PRPN and FDFT1**) have individually been suggested to be involved with the pathogenesis of the disease. **GABBR1** gene encodes a receptor for gamma-aminobutyric acid (GABA), which is the main inhibitory neurotransmitter in the human central nervous system. As observed in this study, the **GABBR1** gene has been previously reported to be down-regulated in AD brains [52]. GABBR1 receptors are prominent in neuronal soma, where NFT formation is known to accumulate. A study examined the immunohistochemical localisation and distribution of GABABR1 protein in the hippocampus of AD subjects and observed a negative correlation with NFT formation and suggested an increase or stable expression of **GBBR1** could contribute to neuronal resistance to the disease process [53].

**GAPDH** gene encodes for a member of the glyceraldehyde-3-phosphate dehydrogenase protein family, which catalyses an important step in the carbohydrate metabolism. GAPDH has been shown to interact with Aβ precursor protein but not cleaved Aβ, and has been proposed to be directly involved in tau aggregation and NFT formation in AD [54]–[56]. The **PRNP** gene encodes for the prion protein, a membrane glycosylphosphatidylinositol-anchored glycoprotein that tends to aggregate into rod-like structures. Mutations in the PRNP gene has been associated with AD and prion protein has also been suggested to be involved in the pathogenesis of AD [57]. **FDFT1** gene encodes a membrane-associated enzyme located at a branch point in the mevalonate pathway, which generate isoprenoids that have been found to be positively correlated with tau pathology [58]. **KLC1** gene encodes for Kinesin Light Chain 1 which transports various cargos such as vesicles, mitochondria, and the Golgi complex along microtubules. An immunoblotting study observed decrease expression of kinesin light chains (KLCs) in the frontal cortex of AD subjects but not in the cerebellum of the same subjects [59]. **TUBA1A** gene encodes for Tublin Alpha 1a, which has been observed to be perturbed in AD [60], and **CLTA** gene encodes for clathrin Light Chain A, which has been observed to be perturbed in AD as well [61]. **COX6C** and **SLC25A5** gene encodes for products which interact with mitochondria and mitochondrial dysfunction in AD has been suggested on numerous occasions [62]–[64].

We identified an additional three AD-specific genes (**UBA1, EIF4H** and **CLDND1**) which were significant DE in all four brain regions. However, the genes were down-regulated in the TL FL and PL but up-regulated in the CB brain region. Ubiquitin-Like Modifier Activating Enzyme 1 (**UBA1**) encodes for a protein that catalyses the first step in ubiquitin conjugation to mark cellular proteins for degradation. Eukaryotic Translation Initiation Factor 4H (**EIF4H**) encodes for a translation initiation factors, which functions to stimulate the initiation of protein synthesis at the level of mRNA utilisation and Claudin Domain Containing 1 (**CLDND1**) is a transmembrane protein of tight junctions found on endothelial cells [65]. **As the cerebellum is the only brain region spared from tangle formation and occasionally from plaque, we suggest these 19 genes (ALDOA, GABBR1, TUBA1A, GAPDH, DNM3, KLC1, COX6C, ACTG1, CLTA, SLC25A5, PRNP, FDFT1, RHOQ, B2M, SPP1, WAC, UBA1, EIF4H and CLDND1)) could potentially be associated with AD histopathology.**

### Translation of proteins perturbed specifically in AD brains

Functional gene set enrichment analysis of the “AD expression profile” revealed more pathways were significantly perturbed in the TL, followed by the FL, PL and CB, which is the general route AD pathology is known to spread through the brain. We originally observed ten biological pathways being enriched across all AD brain regions, which included biological pathways likely to be irrelevant such as the “**digestive system**”. However, when incorporating transcriptomic information from non-AD disorders, we were able to refine the AD expression signature to specific genes perturbed in AD only. This resulted in the enrichment of pathways only involved in the “**metabolism of proteins**”, specifically the translation process which has been previously suggested in be associated with AD on numerous occasions [10] [11] [14] [15] [16] [17]. **We now suggest this may be a biological process specifically disrupted in AD brains, and not BD, HD, MDD, PD or SCZ brains.**

### Previous biological perturbations observed in AD are only associated Temporal Lobe brain region

Previous AD studies have consistently suggested the immune response [10] [11] [12] [13], protein transcription/translation regulation [10] [11] [14] [15] [16] [17], calcium signalling [10] [18] [19], MAPK signalling [16] [7], chemical synapse [18] [7] [19], neurotransmitter pathways [11] [18] [19] and various metabolism pathways [16] [20] [21] [22] [17][11] [23] are disrupted in AD. We observe the same pathways enriched in our meta-analysis; however, only in the TL brain region, a brain region often heavily investigated in AD. Except for “**metabolism of proteins**”, we did not observe any of these pathways significantly enriched across all of the four brain regions, suggesting these pathways observed to be perturbed in previous studies may be tissue-specific rather than disease-specific.

### Interspecies interactions possibly involved in AD

Gene Ontology analysis on the “AD expression profile” identified nine different biological components enriched across all four brain regions. However, when we remove genes perturbed in other neurological or mental health disorders, we only observe four biological components as significantly enriched, and all four were indicative of interspecies interactions. AD brains have a prominent inflammatory component which is characteristic of infection, and many microbes have been implicated in AD, notably herpes simplex virus type 1 (HSV1), Chlamydia pneumonia, and several types of spirochaete [66]. A very recent study also identified common viral species in normal and ageing brains, with an increased human herpesvirus 6A and human herpesvirus 7 in AD brains [67]. Furthermore, Aβ has been suggested to be an antimicrobial peptide and has been shown to protect against fungal and bacterial infections [68]. Thus, the accumulation of Aβ may be part of the brains defence mechanism against infections. Although a controversial theory, we also observe a viral component in AD brains, and as a result of this meta-analysis, further suggest this maybe AD-specific and warrants further investigation.

### Network analysis identifies AD-specific APP UBC and SUMO2 hub genes

Network analysis identified five (**APP, HSP90AA1, UBC, SUMO2** and **RPS2**) significant hub genes specific to AD brain regions. **APP, UBC** and **SUMO2** gene appear as hub genes in multiple brain regions. The **APP** gene encodes for a cell surface receptor transmembrane amyloid precursor protein (APP) that is cleaved by secretases to form a number of peptides. Some of these peptides are secreted and can bind to the acetyltransferase complex APBB1/TIP60 to promote transcriptional activation, while others form the protein basis of the amyloid plaques in AD brains. In addition, two of the peptides are antimicrobial peptides, having been shown to have bacteriocidal and antifungal activities [provided by RefSeq, Aug 2014]. Changes in APP functions have been suggested to play an essential role in the lack of AB clearance, ultimately leading to the formation of plaques [69].

**UBC** (ubiquitin-C) gene encodes for a Polyubiquitin-C protein which is part of the ubiquitin-proteasome system (UPS), the major intracellular protein quality control system in eukaryotic cells. UPS has an immense impact on the amyloidogenic pathway of APP processing that generates Abeta [70]. A recent GWAS study identified **UBC** as a novel LOAD gene, and through network analysis also identified **UBC** as a key hub gene. The study validated their findings in a **UBC** C. elegans model to discover **UBC** knockout accelerated age-related AB toxicity [71]. We also observe the **UBC** gene being down-regulated and as a key hub gene in multiple regions of human AD brains, further providing evidence of its key role in AD.

Small Ubiquitin-Like Modifier 2 (**SUMO2**) gene encodes for a protein that binds to target proteins as part of a post-translational modification system, a process referred to as SUMOylation [72]. However, unlike ubiquitin, which targets proteins for degradation, this protein is involved in a variety of cellular processes, such as nuclear transport, transcriptional regulation, apoptosis, and protein stability [provided by RefSeq, Jul 2008]. Early studies have indicated that the **SUMO** system is likely altered with AD-type pathology, which may impact Aβ levels and tau aggregation [72]. Genetic studies have supported this theory with a GWAS study linking SUMO-related genes to LOAD [73], with further studies showing that the two natively unfolded proteins, tau and α-synuclein, are sumoylated in vitro [74]. We identified **SUMO2** as a significant key hub gene in both the human TL and CB brain region. However, what makes this discovery interesting is that **SUMO2** is significantly up-regulated in the TL, a region where both plaques and tangles can be observed, but significantly down-regulated in the CB, where only plaques have been occasionally observed, but tangles never reported. The up-regulation of **SUMO2** gene may play a vital role in the formation of tangles, and further investigation into this gene is warranted.

### Limitations

Although this study presents novel insights to AD-specific transcriptomic changes in the human brain, limitations to this study must be addressed. Firstly, we meta-analysed a total of 22 AD and 21 non-AD datasets, and many of these datasets lacked necessary experimental processing or basic phenotypic information such as technical batches, RNA integrity numbers (RIN), age, NFT’s, clinical gender, or ethnicity, all of which can have confounding effects. To address this, we incorporated recommended best practices to estimate and correct for both known and hidden batch effects using SVA and COMBAT to ensure data is comparable between experiments and studies. However, this does not guarantee that all technical variation is completely removed.

Secondly, the terminology used to label brain tissue varied across studies, with some reporting a broad region such as the “hippocampus” used in study E-GEOD-48350, while others were very specific to the tissue layer, such as “hippocampus CA3” in study E-GEOD-29378. We, therefore, decided to map all brain tissue as mentioned in each dataset publication to their hierarchical cerebral cortex lobe (TL, FL and PL) and the CB. The mapping procedure was completed using publicly available literature defined knowledge, and we assume tissues within these brain regions are relatively comparable to infer AD-associated histopathological changes.

Thirdly, this study relied on publicly available transcriptomic data, and as previous research has heavily investigated brain regions known to be at the forefront of disease manifestation, this led to unbalanced datasets per brain region in both the AD and non-AD meta-analysis. Subsequently, the AD meta-analysis consisted of 14, 4, 2, and 2 datasets for the TL, FL, PL and CB brain regions respectively, with the PL brain region consisting of only 74 samples (28 AD and 46 controls) in total. In addition, the non-AD meta-analysis lacked expression signatures form all non-AD diseases across all brain regions (except for FL). Nevertheless, the brain regions most affected by each disorder was captured in this study, suggesting we most likely were able to capture key brain transcriptomic changes relating to each disorder. Furthermore, as AD is known to affect all brain regions, albeit not to the same extent, we focus on transcriptomic changes observed across all brain regions that are also not observed in any brain region of the non-AD subjects, ensuring we capture transcriptomic signatures unique to AD brains.

Fourthly, the advances in next sequencing technologies (RNA-seq) which are capable of profiling the whole transcriptome, thus not limited by the pre-defined probes based on known sequencing, would be ideal for disease discoveries. However, AD and mental health studies profiled through RNA-seq is somewhat limited in the public domain, and those that have published DE results are based on small sample numbers, which would fail our selection criteria, such as in [75], [76], [77] and [78]. In addition, these studies lack the same brain regions and mental health disorders covered in this meta-analysis. Nevertheless, we were able to query our genes of interest in the largest known AD RNA-seq web-based database (Agora) which contains DE results from over 2100 human brain samples, however, expression profiling was unavailable for 17/26 genes and DE on the parietal lobe was unavailable. Therefore, this study was unable to comprehensively validate all findings in RNA-seq data.

Finally, we assume the non-AD datasets are comparable through meta-analysis, and by identifying common expression signatures that are not associated with individual disease mechanisms may represent false positives or even a general signature for “brain disorder”. Removing this signature from the AD meta-analysis expression profile may result in transcriptomic changes specific to AD brains, revealing more relevant changes to the underlying disease mechanism rather than general diseases. Under this assumption, we observe more relevant and refined biological enrichment results. For example, we originally observed ten biological pathways enriched across all AD brain regions, including biological pathways such as the “**digestive system**”. However, by refining the AD expression signature by removing genes perturbed in non-AD disorders, only pathways involved in the “**metabolism of proteins**” remain, which has been previously suggested in be associated with AD on numerous occasions [10] [11] [14] [15] [16] [17]. This observation provides strong evidence of our assumption of incorporating non-AD diseases in this study to infer AD-specific changes as valid.

### Conclusion

We present the most extensive human AD brain microarray transcriptomic meta-analysis study to date, incorporating, brain regions both affected and partially spared by AD pathology, and utilise related non-AD disorders to infer AD-specific brain changes. This led to the identification of seven genes specifically perturbed across all AD brain regions and are considered disease-specific, nineteen genes specifically perturbed in AD brains which could play a role in AD neuropathology, and the refinement of GSEA and GO analysis results to identify specific biological pathways and components specific to AD. These AD-specific changes may provide new insights into the disease mechanisms, thus making a significant contribution towards understanding the disease. In addition, two genes (**OGT** and **SOD1**) identified in this study have been specifically silenced in independent mouse models, leading to an accelerated accumulation of hallmark AD pathology and SPCS1 gene was observed to be consistently down-regulated in all AD brain regions based on both microarray and independent RNA-seq data, and has also been nominated as a druggable target for AD treatment. This provides confidence the remaining genes identified in this study may provide new therapeutic targets for AD.

## Acknowledgements

This study presents independent research supported by the NIHR BioResource Centre Maudsley at South London and Maudsley NHS Foundation Trust (SLaM) & Institute of Psychiatry, Psychology and Neuroscience (IoPPN), King’s College London. The views expressed are those of the author(s) and not necessarily those of the NHS, NIHR, Department of Health or King’s College London.

RJBD and SJN are supported by 1. Health Data Research UK, which is funded by the UK Medical Research Council, Engineering and Physical Sciences Research Council, Economic and Social Research Council, Department of Health and Social Care (England), Chief Scientist Office of the Scottish Government Health and Social Care Directorates, Health and Social Care Research and Development Division (Welsh Government), Public Health Agency (Northern Ireland), British Heart Foundation and Wellcome Trust. 2. The National Institute for Health Research University College London Hospitals Biomedical Research Centre

