## Supplementary Text 1 for "A Meta-Analysis of Alzheimer’s Disease Brain Transcriptomic Data"

Supplementary Information 1: Study Outlier process

The following plot was created using the R MetaOmics package’s QC function. This package uses differentially expressed genes (DEGs), co-expression and enriched biological pathways analysis to generate six quantified measures that are used to generate a PCA plot. The direction of each QC measure is juxtaposed on top of the two-dimensional PC subspace using arrows. Datasets in the negative region of the arrows are classed as outliers [1].

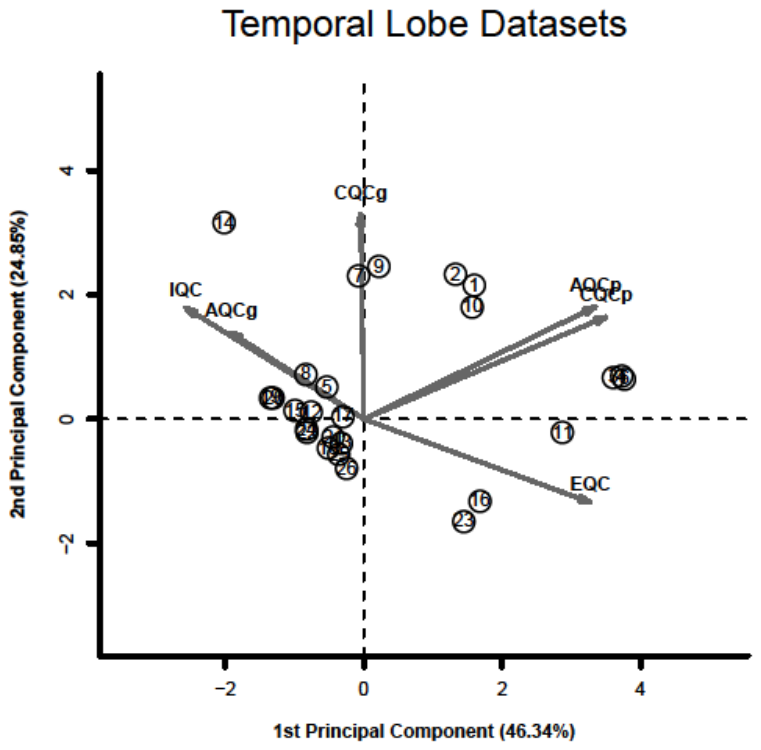

From the 26 temporal lobe datasets plotted, the majority of the AMP MSBB datasets (syn4552659) were in the negative region of the PCA plot arrows, therefore were classed as outliers. The analysis was repeated without the syn4552659 datasets:

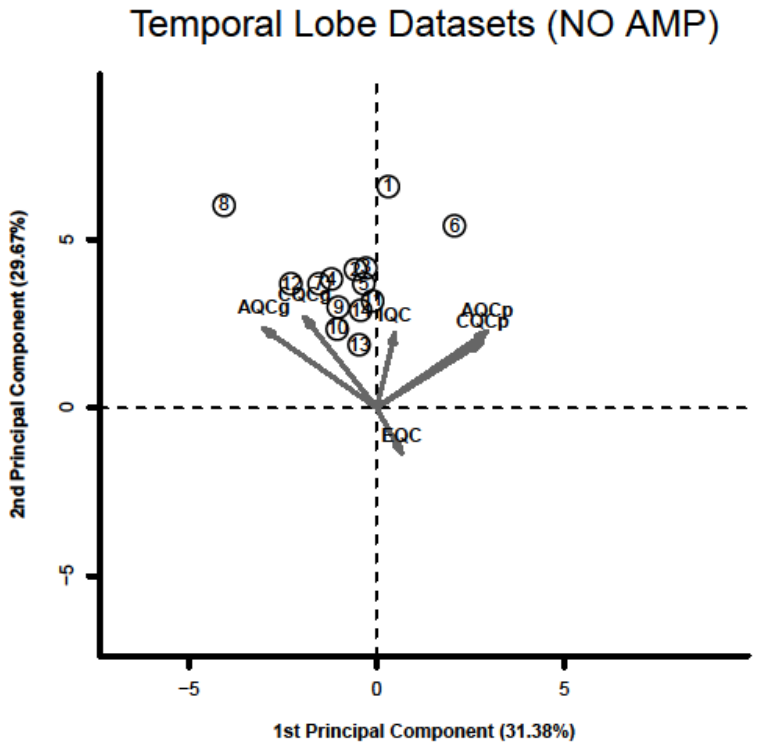

From the remaining 14 datasets, none are classed as clear outliers.
